# Passenger mutations confound phenotypes of SARM1-deficient mice

**DOI:** 10.1101/802702

**Authors:** Melissa B. Uccellini, Susana V. Bardina, Maria Teresa Sánchez-Aparicio, Kris M. White, Ying-Ju Hou, Jean K. Lim, Adolfo García-Sastre

**Affiliations:** Department of Microbiology, New York, NY 10029; Global Health and Emerging Pathogens Institute, New York, NY 10029; Department of Medicine, Division of Infectious Diseases, New York, NY 10029; The Tisch Cancer Institute Icahn School of Medicine at Mount Sinai, New York, NY 10029; Department of Microbiology and Immunology, Weill Medical College of Cornell University, New York, NY 10021

## Abstract

The Toll/IL-1R domain-containing adaptor protein SARM1 is expressed primarily in the brain, where it mediates axonal degeneration. Additional roles for SARM1 in a number of other processes including TLR-signaling, viral infection, chemokine expression, and expression of the proapoptotic protein XAF1 have also been described. Much of the supporting evidence for SARM1 function has been generated by comparing WT C57BL/6 (B6) mice to SARM1-deficient mice backcrossed to the B6 background. Here we show that the *Sarm1* gene lies in a gene-rich region encompassing XAF1, and the MIP and MCP chemokine family loci among other genes. Because gene-targeting of SARM1-deficient strains was done with 129 ES cells and these genes are too close to segregate, they remain 129 in sequence. As this could account for phenotypes attributed to SARM1, we generated new knockout mouse strains on a pure B6 background using CRISPR. Experiments in these new strains confirmed the role of SARM1 in axonal degeneration and susceptibility to WNV infection, but not in susceptibility to VSV or LACV infection, or chemokine or *Xaf1* expression. Notably, the *Xaf1* gene shows sequence variation between B6 and 129, resulting in coding changes and novel splice variants. Given its known role in apoptosis, XAF1 variants may account for some phenotypes described in previously made SARM1-deficient strains. RNAseq in the new strains reveal changes in the mitochondrial electron transport chain and ribosomal proteins, suggesting possible downstream targets of SARM1. Re-evaluation of described phenotypes in these new strains will be critical for defining the function of SARM1.

## Introduction

Sterile alpha and TIR motif containing 1 (SARM1) is an intracellular protein that is highly expressed in the brain, and is comprised of a C-terminal Toll-interleukin receptor (TIR) domain, 2 central sterile alpha motif (SAM) domains, and an N-terminal region containing multiple armadillio repeat motifs (ARMs) (1). SARM1 is essential in Wallerian degeneration - a neuronal cell death program involving MAPK signaling, influx of calcium, and proteolysis of structural proteins resulting in axonal degeneration distal to the site of injury (2, 3). Although the mechanism is not fully elucidated, SARM1 appears to be the master executioner in this cascade (4). Mechanistic and structural studies suggest that the SARM1 TIR domain possesses intrinsic NAD+ cleavage activity (5-7), which is regulated by JNK-mediated phosphorylation of Ser-548 leading to inhibition of mitochondrial respiration (8).

Because of the presence of the TIR domain, it was originally postulated that SARM1 would function in TLR signaling similar to the other cytosolic TIR-domain containing proteins MYD88, MAL, TRIF, and TRAM. In addition, the *C.elegans* and *Drosophila* orthologs *tir-1* and *dSARM (ect-4)* appear to have roles in immunity (9-11). However, unlike the other four adaptor proteins, overexpression of SARM1 did not lead to NF-κB or IRF3 activation, but rather inhibited TLR signaling (12). Overexpression studies have supported a role for SARM1 in suppressing TLR responses, however studies in knockout mice have not (1). Importantly, the SARM1 TIR domain appears to be evolutionarily ancestral to the mammalian TLR adaptors because of its closer homology to bacterial TIR domains, suggesting that it may not function as a TLR adapter (13, 14).

SARM1 also appears to play a role in susceptibility to infections of the CNS. Two knockout strains for SARM1 have been generated, one in the Ding lab here referred to as *Sarm1*^*AD*^ (1) and one in the Diamond lab here referred to as *Sarm1*^*MSD*^ (15). *Sarm1*^*MSD*^ mice are more susceptible to West Nile virus infection (WNV), and produce less TNF-α (15). In contrast, *Sarm*^*MSD*^ mice are protected from lethal La Crosse virus infection (LACV) (16). Our previous studies found that *Sarm1*^*AD*^ mice were also protected from lethal Vesicular Stomatitis virus (VSV) infection, and produced less cytokines and chemokines in the brain (17). A role for SARM1 has only been shown for infections in the CNS – we did not find differences in the susceptibility of *Sarm1*^*AD*^ mice to *M.tuberculosis, L. monocytogenes*, or influenza virus infection (17). When *Sarm1*^*AD*^ macrophages were examined in response to a variety of TLR ligands no differences were found in the production of TNF-α or CCL2 (1). However, SARM1 was reported to regulate CCL5 production in *Sarm1*^*AD*^ macrophages. This defect was specific to CCL5, occurred in response to TLR and non-TLR stimuli, did not involve known signaling intermediates, but was associated with recruitment of RNA pol II and transcription factors to the CCL5 locus (18). A recent report also described both positive and negative roles for SARM1 in inflammasome activation in *Sarm1*^*AD*^ mice, whereby SARM1 positively regulates pyroptosis but negatively regulates IL-1β secretion (19).

We previously reported upregulation of *Xaf1* transcripts in the brains of uninfected and VSV-infected *Sarm1*^*AD*^ mice compared to WT mice (17). Zhu et al recently described a similar phenotype in *Sarm*^*MSD*^ mice, and reported that SARM1 modulates *Xaf1* transcript expression and caspase-mediated cell death (20). X-linked inhibitor of apoptosis (XIAP)-associated factor (XAF1) is a proapoptotic IFN-stimulated gene that is epigenetically silenced in a broad range of human tumors. XAF1 appears to induce apoptosis through a variety of mechanisms including binding and inhibiting XIAP (21), and binding p53 displacing MDM2 leading to cell death (22). Several isoforms of *Xaf1* have been described, including full-length and truncated forms. Full-length isoforms are frequently downregulated in human tumors, while truncated isoforms are upregulated. Importantly, short forms have been reported to have dominant negative effects (22, 23).

## Results

### Macrophages derived from Sarm1^AD^ mice are defective in the production of Ccl3, Ccl4, and Ccl5

We stimulated bone marrow-derived macrophages with TLR ligands, or infected with viruses known to activate the RLR sensing pathway and measured cytokine and chemokine production by ELISA. For this purpose we compared WT C57BL/6J (B6) mice to SARM1-deficient mice generated in the Ding lab and backcrossed 10 times to the B6 background, here referred to as *Sarm1*^*AD*^ (see Table I for background details of the mice used in this study). We found that while TNF-α and IFN-α production were normal in *Sarm1*^*AD*^ macrophages, CCL3 production was defective in response to all stimuli tested (Fig 1A). We next asked if the defect in chemokine production occurred at the transcriptional level. *Sarm1*^*AD*^ macrophages showed defects in the production of *Ccl3, Ccl4*, and *Ccl5* mRNA in response to LPS stimulation at a number of time points, but no defects in the production of *Il1b* or *Ifnb1* (Fig 1B, top), similar to results reported for *Ccl5* (18). Given that we saw defects in chemokine production in response to a variety of TLR stimuli, we next asked if signaling in response to TNF-α, which does not use the TLR adaptor proteins MYD88 or TRIF, was defective in *Sarm1*^*AD*^ macrophages. *Sarm1*^*AD*^ macrophages again showed defects in the production of *Ccl3, Ccl4*, and *Ccl5* mRNA, but not *Il1b* or *Ifnb1* (Fig 1B, bottom). This suggested that the defect in chemokine production in *Sarm1*^*AD*^ macrophages was not specific to the TLR signaling pathway.

**Table I.** Summary of *Sarm1* mouse lines and phenotypes

| Allele | Ref | Neo | Congenic interval | Genetic background | Axonal degeneration | VSV | LACV | WNV | Chemokines |
| --- | --- | --- | --- | --- | --- | --- | --- | --- | --- |
| *Sarm1<sup>AD</sup>* | 1 | Y | 129 | 99.5% B6 | protected | ↓ | WT (original strain) | WT | ↓ |
| *Sarm1<sup>MSD</sup>* | 15 | N | 129 | 94.6% B6 | * | ND | ↓(1978 strain) | ↑ | ND |
| *Sarm1<sup>AGS3</sup>* | this study | N | None | 100% B6 | protected | WT | WT (original strain) | ↑ | WT |
| *Sarm1<sup>AGS12</sup>* | this study | N | None | 100% B6 | ND | WT | WT (original strain) | ND | WT |

ND=not done, ↓= decreased susceptibility, ↑= increased susceptibility \*Protection from axonal degeneration has been shown in many studies, but the strain is often not specified

**FIGURE 1.**
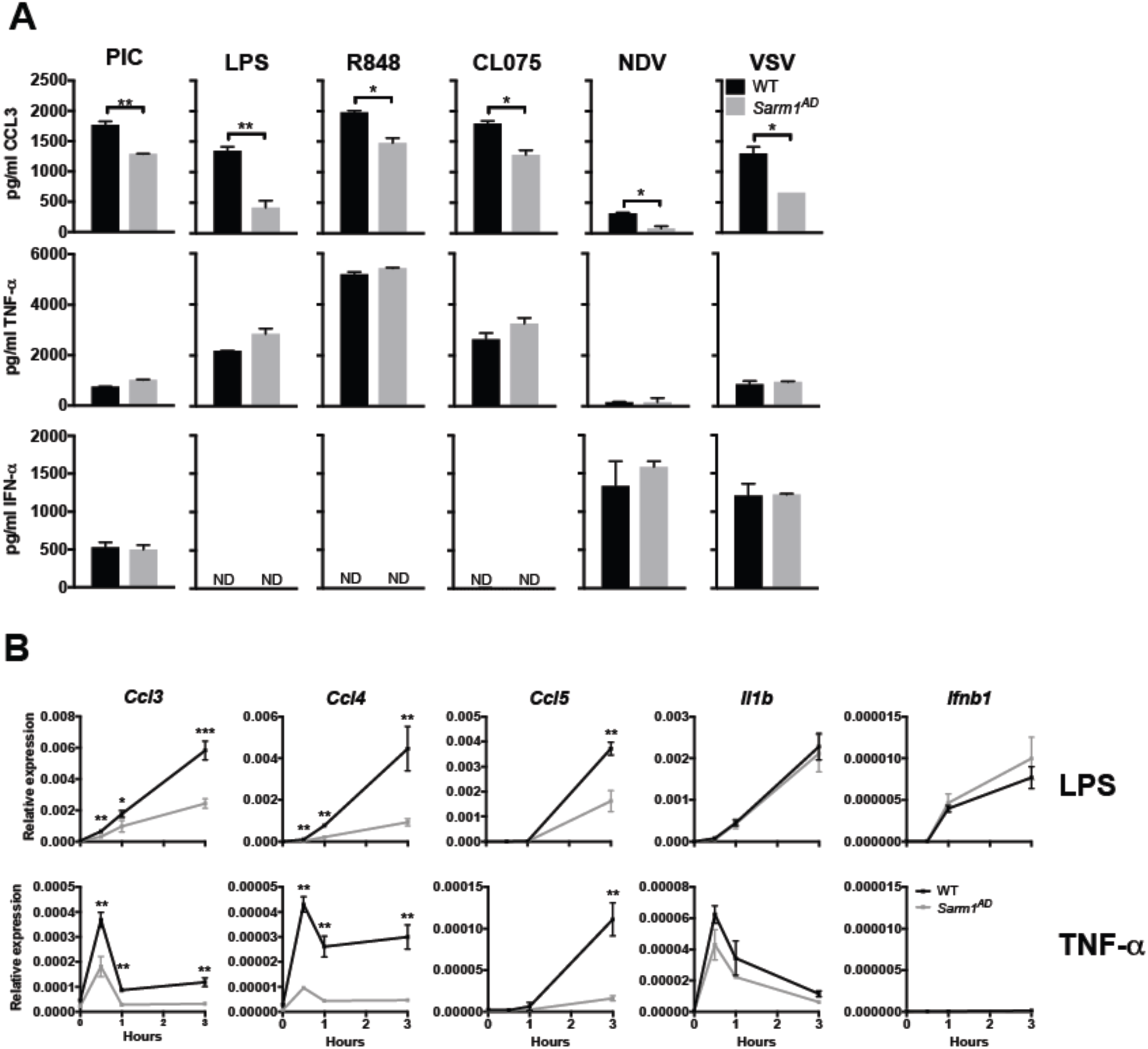
Macrophages from *Sarm1*^*AD*^ mice have a defect in the production of *Ccl3, Ccl4*, and *Ccl5*. (**A**) WT and *Sarm1*^*AD*^ macrophages were stimulated with 100 μg/ml PIC, 5 μg/ml LPS, 0.01 μg/ml R848, 10 μg/ml CL075, or NDV or VSV at an MOI of 5 for 24 hrs. and cytokine production was measured by ELISA. (**B**) WT and *Sarm1*^*AD*^ macrophages were stimulated with 1 μg/ml LPS or TNF-α and cytokine production was measured by qPCR at the indicated time points. Graphs show mean+/-SD for triplicate biological replicates and are representative of 3 experiments. *p<0.05 and * *p<0.01 (unpaired t test).

### Macrophages derived from Sarm1^AD^ mice show normal signaling responses

We saw defects in the production of chemokines in *Sarm1*^*AD*^ macrophages in response to both LPS and TNF-α stimulation, suggesting that SARM1 does not function at the level of the TLR-adapter proteins MYD88 or TRIF. However, both LPS and TNF-α signaling activate the NF-κB and MAPK signaling pathways (24, 25). We therefore examined activation of these pathways in *Sarm1*^*AD*^ macrophages by western blot. No differences were observed in the degradation of IκBα, or the phosphorylation of JNK, ERK, or p38 in response to either LPS or TNF-α stimulation, suggesting that SARM1 does not regulate induction of the NF-κB or MAPK pathways (Fig 2A and B). LPS also activates PI3 kinase signaling resulting in phosphorylation of Akt (26), however no differences in p-Akt levels were observed in *Sarm1*^*AD*^ macrophages in response to LPS (Fig 2C). In addition, PLCγ-2 and intracellular calcium are required for TLR4 endocytosis in response to LPS (27). However, we again saw no differences in intracellular Ca^2+^ flux in *Sarm1*^*AD*^ macrophages in response to LPS or ATP stimulation (Fig 2D).

**FIGURE 2.**
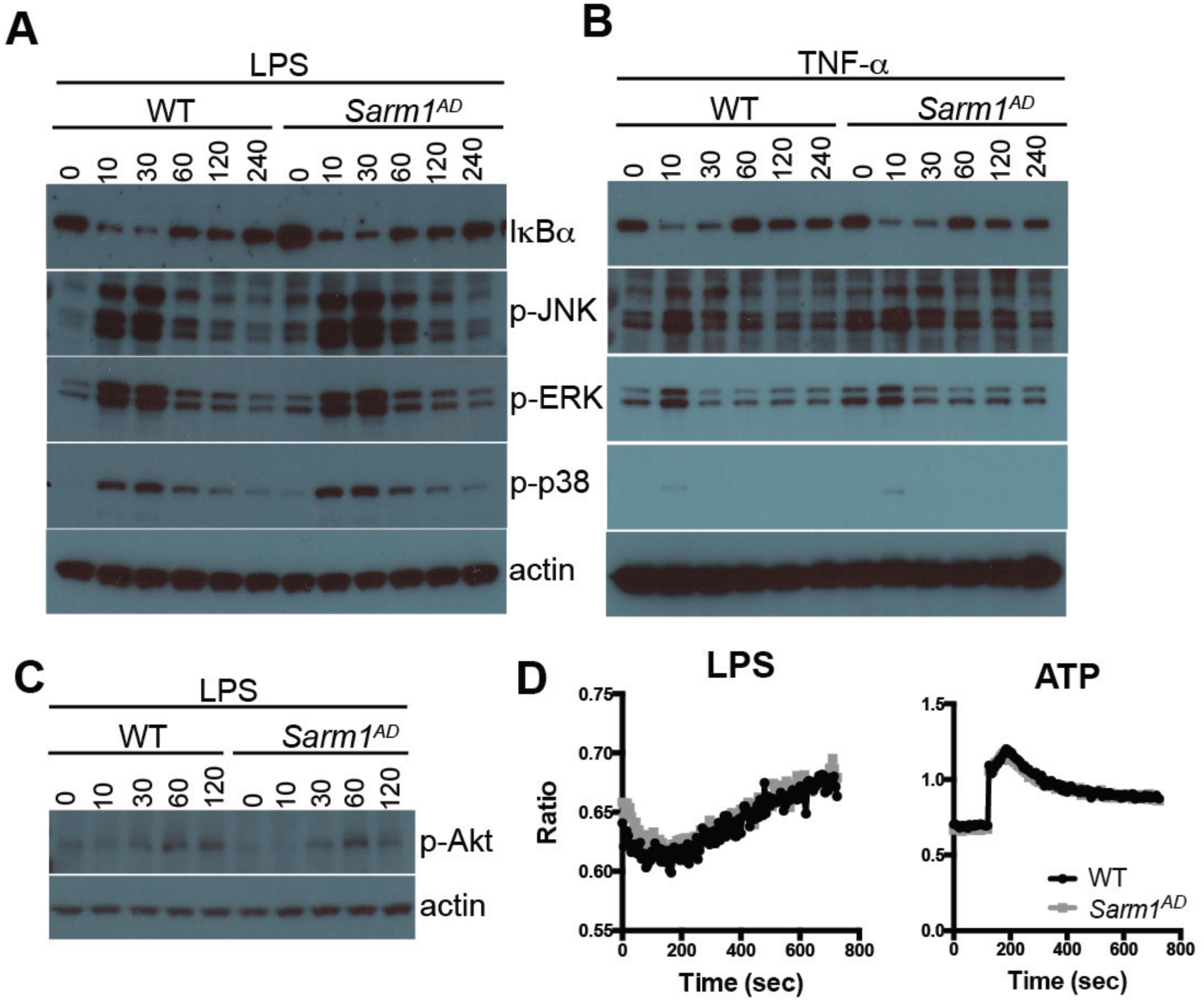
Macrophages from *Sarm1*^*AD*^ mice display normal signaling responses. WT and *Sarm1*^*-/-*^ macrophages were stimulated with 10 ng/ml LPS (**A** and **C**) or TNF-α (**B**) for the indicated number of minutes and signaling responses were measured by western blot. (**D**) WT and *Sarm1*^*-/-*^ macrophages were stimulated with 100 ng/ml LPS or 1 mM ATP and calcium flux was measured by fura-2-AM fluorescence. Data is representative of 3 experiments.

### The MIP and MCP chemokine family loci are within the Sarm1 129 congenic locus

Given that we saw defects only in *Ccl3, Ccl4*, and *Ccl5* production but not in other cytokines, that the defects occurred in response to a wide variety of stimuli, and that no defects in the induction pathways for these cytokines could be found – we considered the possibility that the observed defect was due to the genetic background of the knockout mouse rather than lack of SARM1 expression. The *Sarm1*^*AD*^ strain was made by replacing exons 3-6 with a neomycin resistance gene in reverse orientation in 129 ES cells, before backcrossing 10 times to the B6 background (1). The *Ccl3, Ccl4*, and *Ccl5* genes and the *Sarm1* gene are both located on mouse chromosome 11, and are separated by only ∼5 Mb (Fig 3A). Despite backcrossing 10 times, the probability of a region of 5 cM (∼6.75 Mb for chromosome 11(28)) of 129 genetic material flanking both sides of the knockout gene is 0.63, making it likely that the chemokine locus in *Sarm1*^*AD*^ mice is of 129 origin. In order to check the genetic background of genes proximal to *Sarm1*, we sequenced two SNPs in the *Ccl5* gene that differ between the 129 and B6 strains, which confirmed that the *Ccl5* locus of the *Sarm1*^*AD*^ strain is derived from the 129 strain (Fig 3B).

**FIGURE 3.**
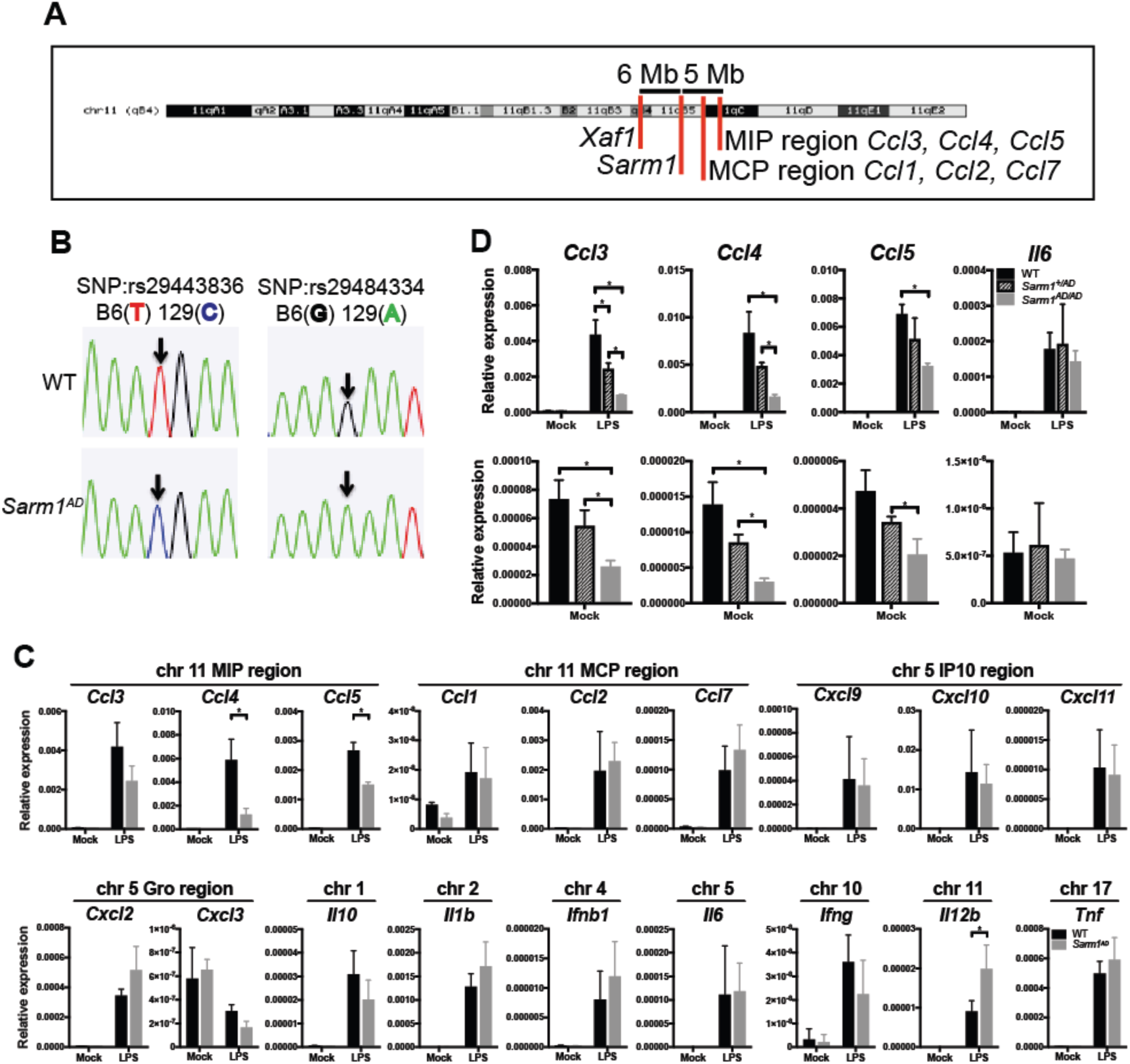
*Ccl3, Ccl4, Ccl5* and *Xaf1* are within the *Sarm1*^*AD*^ 129 congenic locus. (**A**) Chromosomal location of the *Sarm1* gene, chemokine locus, and *Xaf1* gene (UCSC genome browser). (**B**) Sequence analysis of SNPs in the *Ccl5* gene of WT and *Sarm1*^*AD*^ mice. (**C**) WT and *Sarm1*^*AD*^ macrophages were stimulated with 10 ng/ml LPS for 3 hrs. and cytokine production was measured by qPCR. (**D**) WT, *Sarm1*^*+/AD*^, and *Sarm1*^*AD/AD*^ macrophages were stimulated as in C (bottom graph shows the same data as the top on a different scale). C and D show mean+/-SD for triplicate biological replicates and are representative of 2 experiments. *p<0.05 (unpaired t test).

We next asked whether the production of other cytokines and chemokines located on different chromosomes was different between WT and *Sarm1*^*AD*^ macrophages. We again saw differences in the production of *Ccl3, Ccl4*, and *Ccl5* mRNA, but we failed to find significant differences between other cytokines or chemokines in different chromosomal locations (Fig 3C). The MCP chemokine region falls between the *Sarm1* gene and the MIP chemokine region, and is therefore of 129 genetic origin, however no differences in the induction of *Ccl1, Ccl2*, or *Ccl7* were observed. *Il12b*, which is also located on chromosome 11, showed increased production in the *Sarm1*^*AD*^ strain. In addition to induced conditions (Fig 3D, top), we also observed differences in the basal expression of *Ccl3, Ccl4*, and *Ccl5* mRNA between WT, *Sarm1*^*+/AD*^, and *Sarm1*^*AD/AD*^ macrophages in the absence of stimulation (Fig 3D, bottom), supporting an intrinsic difference between the strains.

### SARM1 knockdown and overexpression fail to regulate Ccl3, Ccl4, and Ccl5 levels

We next examined the role of SARM1 expression on chemokine production in a cell line, lacking the confounding genetic background of the *Sarm1*^*AD*^ mouse strain. We first examined *Sarm1* expression in the mouse macrophage cell line RAW264.7 expressing a control V5 epitope tag (RAW-V5). We found very low levels of *Sarm1* mRNA expression, making knockdown efficiency difficult to access (Fig 4A, left). This is in agreement with reports suggesting very low or no expression in mouse macrophages (1, 15). Upon treatment with LPS, no differences in *Ccl4* induction were found with knockdown (Fig 4A, right). In order to determine knockdown efficiency, we repeated the experiment in RAW264.7 cells overexpressing V5-tagged SARM1 (RAW-SARM1-V5). Under these conditions, *Sarm1* mRNA was detectable, and siSARM1-1 and siSARM1-3 reduced transcript expression by 10x and 7x, respectively, confirming knockdown (Fig 4B, left). Western blot for Sarm1-V5 expression revealed siSARM1-1 and siSARM1-3 reduced protein levels by 40% and 30%, respectively (Fig 4C and S1). The low knockdown efficiency is likely due to high SARM1 expression from the CMV promoter, but nonetheless confirms the efficacy of the siRNAs. However, upon LPS stimulation, again no differences in *Ccl4* mRNA induction were detectable in RAW-SARM1-V5 cells (Fig 4B, right). We next performed knockdown in macrophages from WT and *Sarm1*^*AD*^ mice. We were unable to detect *Sarm1* mRNA expression in macrophages, and no reliable antibodies are available (1, 15, 17, 18), so we could not access knockdown efficiency. We again found that basal levels of *Ccl4* mRNA were reduced in *Sarm1*^*AD*^ macrophages compared to WT macrophages, however siRNA treatment of WT macrophages failed to downregulate *Ccl4* levels (Fig 4D). Lastly, we determined whether overexpression of SARM1 in RAW cells modulated chemokine induction in response to LPS. As shown in Figure 4E, no differences in chemokine levels were observed upon overexpression of SARM1. The limited chemokine defects, lack of signaling defects, and lack of support from knockdown or overexpression, as well as the close proximity of the *Ccl3, Ccl4*, and *Ccl5* genes to the *Sarm1* gene makes it likely that the congenic interval rather than SARM1 protein expression contributes to differences in basal and induced levels of *Ccl3, Ccl4*, and *Ccl5* between WT and *Sarm1*^*AD*^ mice.

**FIGURE 4.**
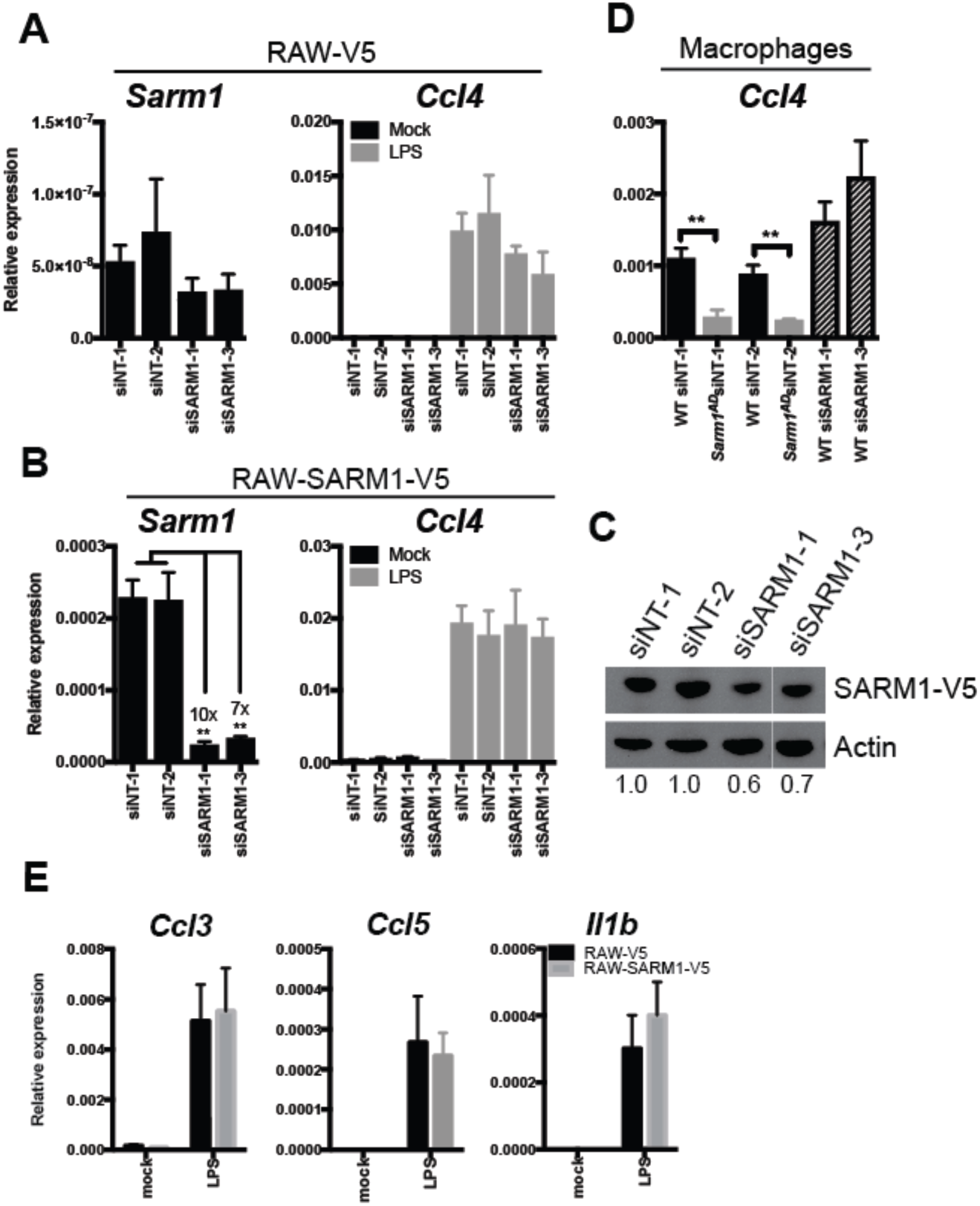
SARM1 knockdown and overexpression do not modulate chemokine production. (**A**) RAW-V5 cells were treated with *Sarm1* siRNAs and *Sarm1* knockdown efficiency was measured by qPCR (left) or *Ccl4* expression was measured after treatment with 10 ng/ml LPS for 3 hrs. (right). (**B**) RAW-SARM1-V5 cells treated as in A. (**C**) Western blot of SARM1 expression in RAW-SARM1-V5 cells treated with *Sarm1* siRNAs. (**D**) Basal expression of *Ccl4* in WT and *Sarm1*^*AD*^ macrophages by qPCR after *Sarm1* siRNA knockdown. (**E**) RAW-V5 and RAW-SARM1-V5 cells were treated with 10 ng/ml LPS and cytokine production was measured at 3 hrs. by qPCR. Graphs show mean+/-SD of triplicate biological samples and are representative of 2 experiments. * *p<0.01 (unpaired t test).

### Sarm1 CRISPR knockout mice on a pure B6 background show no macrophage chemokine defects

In order to formally exclude a role for SARM1 in chemokine induction, we generated new knockout mouse strains using CRISPR-mediated genome engineering on a pure B6 background. A high-scoring guide sequence that was unlikely to produce off-target cleavage was located in exon 1 of the *Sarm1* gene (29). This guide sequence was cloned into the pSpCas9(BB)-2A-GFP vector and injected into one-cell stage C57BL/6J embryos. Resulting pups were characterized at the *Sarm1* locus, as well as at potential off-target sites. Two knockout alleles were generated using this approach, termed *Sarm1*^*AGS3*^ and *Sarm1*^*AGS12*^. The *Sarm1*^*AGS3*^ allele is a 62 b.p. deletion resulting in a frameshift and a 38 a.a. product; the *Sarm1*^*AGS12*^ allele is a 13 b.p. deletion resulting in a frameshift and a 74 a.a. product (Table II and Fig S2A). The 62 b.p. deletion in the *Sarm1*^*AGS3*^ allele was evident by PCR of *Sarm1* genomic DNA (Fig S2B, left). The 13 b.p. deletion in the *Sarm1*^*AGS12*^ allele was too small to be detected on an agarose gel, but was detected using the Surveyor Nuclease assay (S2B, right). The guide sequence used for *Sarm1* cleavage was high scoring and no potential off-target sites were present with less than 4 mismatches, making CRISPR cleavage at off-target sites unlikely (30). Nonetheless, we tested 5 potential off-target sites located in exonic regions that could potentially affect these genes. We did not detect cleavage events at any of these sites as determined by the Surveyor Nuclease assay (Fig S2C).

**Table II.**
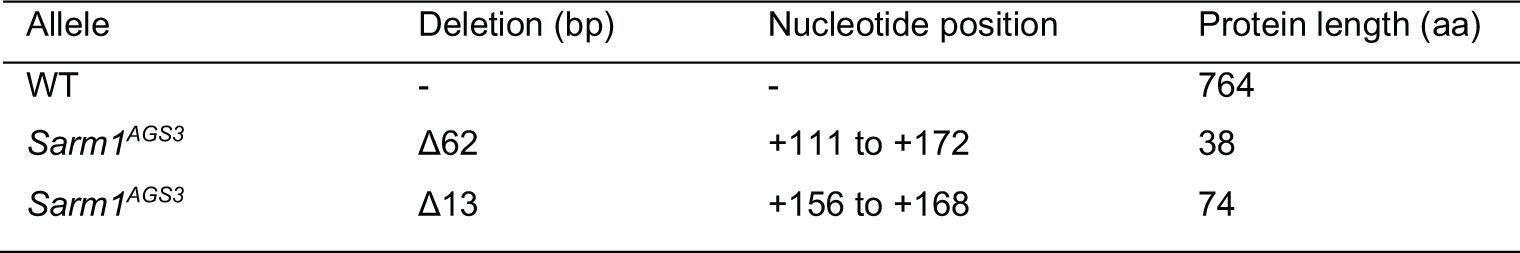
*Sarm1* Alleles

| Allele | Deletion (bp) | Nucleotide position | Protein length (aa) |
| --- | --- | --- | --- |
| WT | - | - | 764 |
| *Sarm1*<sup>AGS3</sup> | Δ62 | +111 to +172 | 38 |
| *Sarm1*<sup>AGS12</sup> | Δ13 | +156 to +168 | 74 |

The *Sarm1*^*AGS3*^ and *Sarm1*^*AGS12*^ lines were breed to homozygosity creating two new *Sarm1* knockout strains. We compared responses of macrophages derived from WT, the original *Sarm1*^*AD*^ line, and the *Sarm1*^*AGS3*^ line. As expected, the *Sarm1*^*AD*^ macrophages showed defects in the production of *Ccl3, Ccl4*, and *Ccl5* mRNA in response to LPS (Fig 5A, top) or TNF-α (Fig 5A, bottom). However, the *Sarm1*^*AGS3*^ line showed responses comparable to WT. The *Sarm1*^*AGS12*^ line also showed *Ccl3, Ccl4*, and *Ccl5* responses comparable to WT in response to TNF-α (Fig 5B). This shows that defects in the production of chemokines in the original *Sarm1*^*AD*^ macrophages were due to background effects, and not SARM1 protein expression.

**FIGURE 5.**
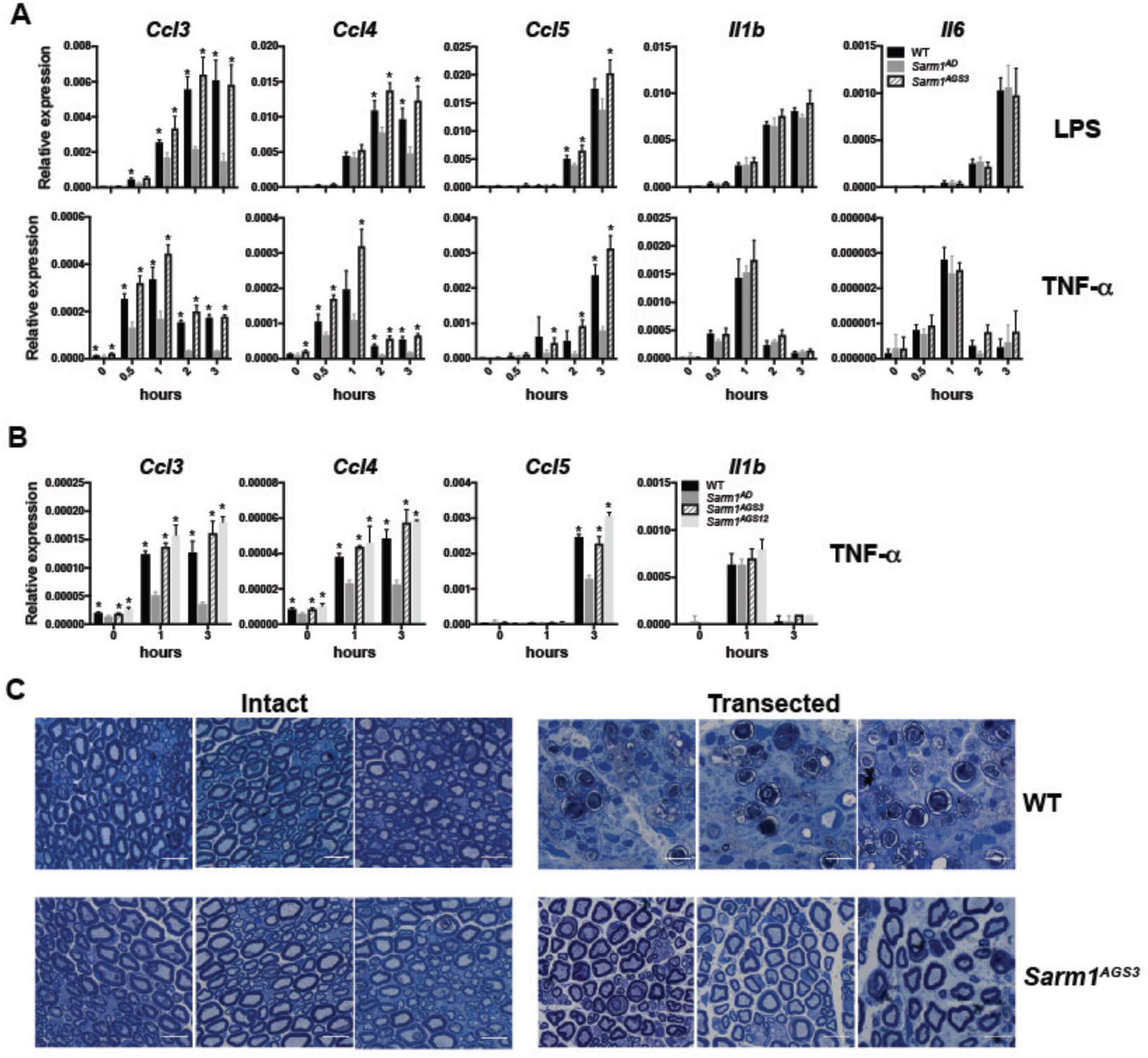
*Sarm1* CRISPR knockout mice on a pure B6 background show normal chemokine production, but are protected from axonal degeneration. (**A**) WT, *Sarm1*^*AD*^, and *Sarm1*^*AGS3*^ macrophages were stimulated with 10 ng/ml LPS or TNF-α and cytokine production was measured at the indicated time points by qPCR. (**B**) WT, *Sarm1*^*AD*^, *Sarm1*^*AGS3*^, and *Sarm1*^*AGS12*^ macrophages were stimulated with 10 ng/ml TNF-α as in A. (**C**) Toluidine Blue staining of sciatic nerves from untransected (left) and transected (right) WT and *Sarm1*^*AGS3*^ mice 14 days post-transection. Scale bar 10μm. Graphs show mean+/-SD of triplicate biological samples and are representative of 3 experiments. *p<0.05 (unpaired t test).

### Sarm1 CRISPR knockout mice are protected from axonal degeneration

We have been unable to detect the expression of a SARM1-specific band by western blot using a number of commercial antibodies and western blotting conditions (not shown). We therefore sought to confirm knockout of SARM1 protein expression functionally in an axonal degeneration assay. For this purpose, we performed sciatic nerve transections of the right hindlimb in WT and *Sarm1*^*AGS3*^ mice. 14 days following transection, WT mice showed breakdown of the axon and myelin sheath, while *Sarm1*^*AGS3*^ mice showed remarkable protection (Fig 5C) as described previously in the *Sarm1*^*AD*^ strain (2). This confirms a role for SARM1 in axonal degeneration, and functional knockout of SARM1 in the *Sarm1*^*AGS3*^ line.

### Viral phenotypes of Sarm1 CRISPR mice

We had previously reported that *Sarm1*^*AD*^ mice are resistant to lethal encephalitic disease caused by VSV infection (17). In order to determine if this was a true function of SARM1, we infected *Sarm1*^*AGS3*^ and *Sarm1*^*AGS12*^ mice with VSV and monitored survival. As shown in Figure 6A, *Sarm1*^*AD*^ mice, but not *Sarm1*^*AGS3*^ or *Sarm1*^*AGS12*^ mice were protected from VSV, suggesting that SARM1 does not play a role in VSV infection. Our reported defects in cytokine and chemokine production in the brain of VSV-infected mice were also due to background effects and not SARM1 protein (Fig 6B). An independent line of SARM1-deficient mice (referred to here as *Sarm1*^*MSD*^) was generated in the Diamond lab also on the 129 background but lacking the neomycin cassette. These mice showed increased susceptibility to WNV infection (15). When *Sarm1*^*AGS3*^ mice were infected with WNV, they were more susceptible than WT mice (Fig 6C) confirming a role for SARM1 in WNV infection in agreement with the Diamond study. Surprisingly, *Sarm1*^*AD*^ mice showed similar susceptibility to WT mice to WNV infection (Fig 6C and Table I), suggesting that background effects in *Sarm1*^*AD*^ mice may have compensated for the impact of SARM1-deficiency on susceptibility to WNV infection. *Sarm1*^*MSD*^ mice were also reported to be protected from LACV infection (16). When *Sarm1*^*AD*^, *Sarm1*^*AGS3*^, and *Sarm1*^*AGS12*^ mice were infected with LACV, all strains showed similar susceptibility to WT mice, suggesting that SARM1 also does not play a role in susceptibility to LACV infection (Fig 6D).

**FIGURE 6.**
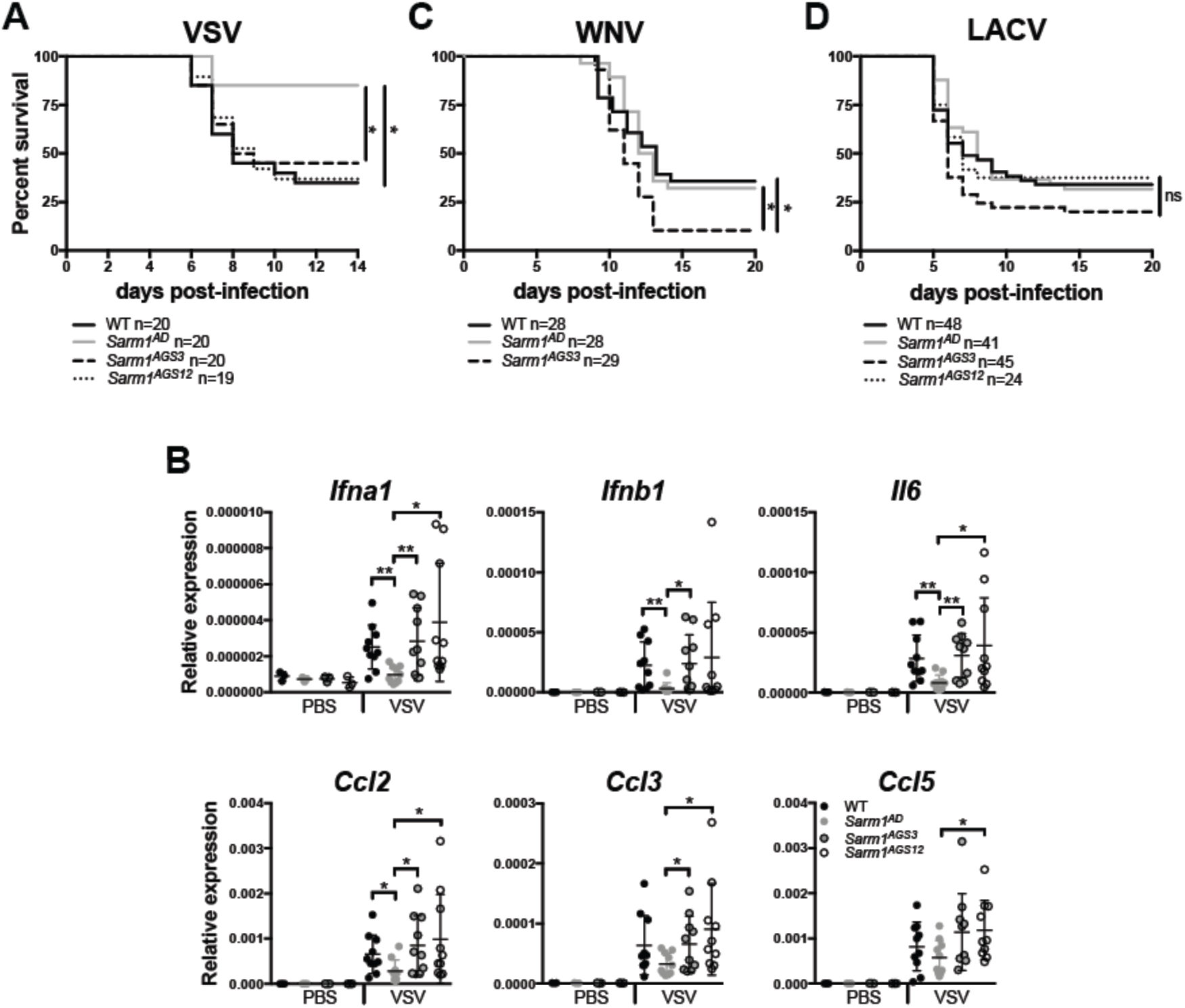
Viral phenotypes of *Sarm1* CRISPR knockout mice. (**A**) WT, *Sarm1*^*AD*^, *Sarm1*^*AGS3*^, and *Sarm1*^*AGS12*^ mice were infected intranasally with 10^7^ pfu of VSV and survival was measured. (**B**) Mice were infected as in C, and chemokine production in the brain was measured by qPCR at day 6 post-infection. (**C**) WT, *Sarm1*^*AD*^, and *Sarm1*^*AGS3*^ mice were infected with 10^2^ FFU of WNV-NY99 via footpad injection and survival was measured. **(D)** WT, *Sarm1*^*AD*^, and *Sarm1*^*AGS3*^, and *Sarm1*^*AGS12*^ mice were infected intraperitoneally with 10^3^ pfu of LACV original strain and survival was measured. A, C, and D show combined results of 2 experiments with similar results, B shows mean+/-SD for n=3 (PBS) and n=10 (VSV) and are representative of 3 experiments. *p<0.05 log-rank test (A, C, D) unpaired t test (B).

The *Sarm1*^*AD*^ mice used in this study were backcrossed 10 times to the B6 background; *Sarm1*^*MSD*^ were reported to be backcrossed to the B6 background, however the extent of backcrossing was not reported. In order to determine the precise backgrounds of the two strains, we performed a 384 panel SNP analysis. The *Sarm1*^*AD*^ mice were 99.5% B6, while the *Sarm1*^*MSD*^ mice were 94.6% B6. The *Sarm1*^*AD*^ mice were found to differ from B6 at the expected location on chromosome 11 and one other region on chromosome 10. The *Sarm1*^*MSD*^ mice were found to differ from B6 at multiple locations including large portions of chromosome 10 and 11 (Table S1), which may account for the different phenotypes observed with the two strains. It should be noted that the precise genetic background of the strains used in different labs and studies will likely differ depending on the extent of backcrossing done in individual labs.

**Table S1.** SNP analysis of *Sarm1*^*AD*^ and *Sarm1*^*MSD*^ mice

| SNP# | Chromosome | NCBI Assembly bp | B6J | *Sarm1<sup>AD</sup>* | *Sarm1<sup>MSD</sup>* |
| --- | --- | --- | --- | --- | --- |
| 11 | 01-11 | 75525463 | V V |  | F F |
| 21 | 01-21 | 144953207 | F F |  | V V |
| 115 | 05-15 | 104143745 | V V |  | V F |
| 180 | 08-14 | 88318808 | F F |  | V V |
| 205 | 10-01 | 7775347 | F F |  | V F |
| 207 | 10-03 | 20325683 | F F |  | V F |
| 209 | 10-05 | 38965551 | V V |  | V F |
| 211 | 10-07 | 48685723 | F F |  | V F |
| 213 | 10-09 | 67238174 | F F | V F | V V |
| 225 | 11-01 | 9552730 | F F |  | V F |
| 229 | 11-05 | 37252460 | V V |  | F F |
| 230 | 11-06 | 45819381 | F F |  | V V |
| 231 | 11-07 | 54267162 | V V |  | F F |
| 233 | 11-09 | 65095597 | V V |  | F F |
| 235 | 11-11 | 75647055 | F F | V V | V V |
| 248 | 12-06 | 34055903 | F F |  | V F |
| 260 | 12-18 | 114730692 | F F |  | V F |
| 275 | 13-15 | 102573089 | F F |  | V F |
| 277 | 13-17 | 115537300 | F F |  | V F |
| 290 | 14-12 | 86216087 | V V |  | V F |
| 292 | 14-14 | 101001947 | V V |  | V F |
| 297 | 15-01 | 10575512 | F F |  | V F |
| 299 | 15-03 | 22519205 | F F |  | V F |
| 301 | 15-05 | 35126500 | F F |  | V V |
| 329 | 17-03 | 23972821 | F F |  | V F |
| 331 | 17-05 | 33380773 | F F |  | V F |
| 349 | 18-09 | 58557879 | F F |  | V F |
| 356 | 19-02 | 17770192 | F F |  | V F |
| 362 | 19-08 | 59241076 | V V |  | V F |

F=FAM probe
V=VIC probe

### Xaf1 expression differences are due to sequence and isoform polymorphism between B6 and 129

Significant differences in transcript levels of *Xaf1*, a proapoptotic protein, were reported by us in the original *Sarm1*^*AD*^ strain both in the presence and absence of VSV infection, and others (20) in the *Sarm1*^*MSD*^ strain both in the presence and absence of prion infection. Additionally, *Xaf1* was the most highly upregulated transcript in SARM1-deficient mice compared to WT mice in both studies. Two curated protein-coding transcripts for *Xaf1* have been described in mouse (Fig 7A), as well as a number of predicted transcripts. Isoform 1 contains exons 1-6 and isoform 2 contains exons 1, 2, 5, and 6. The *Xaf1* gene is also located in close proximity to the *Sarm1* gene on chromosome 11 (Fig 3A). Alignment of RNAseq reads from the *Sarm1*^*AD*^ strain to the B6 reference genome showed a number of nucleotide differences (Fig 7B – indicated by colored lines), and the *Sarm1*^*AD*^ consensus sequence matched the reported sequence for 129. The nucleotide differences in exons 4 and 5 result in 4 amino acid substitutions (Fig 7E). The 129 sequence has a gap in the alignment at the 3’ end of exon 6, which is the result of a 248 bp deletion, and a large peak in the 3’ UTR that is not present in B6. The deletion spans the B6 stop codon and 2 polyadenylation sites, which likely results in a transcript that terminates much later in 129, potentially effecting transcript stability. The 129 transcript uses an alternative stop codon located after the deletion, resulting in truncation of the last 3 amino acids at the C-terminus of the protein (Fig 7E).

**FIGURE 7.**
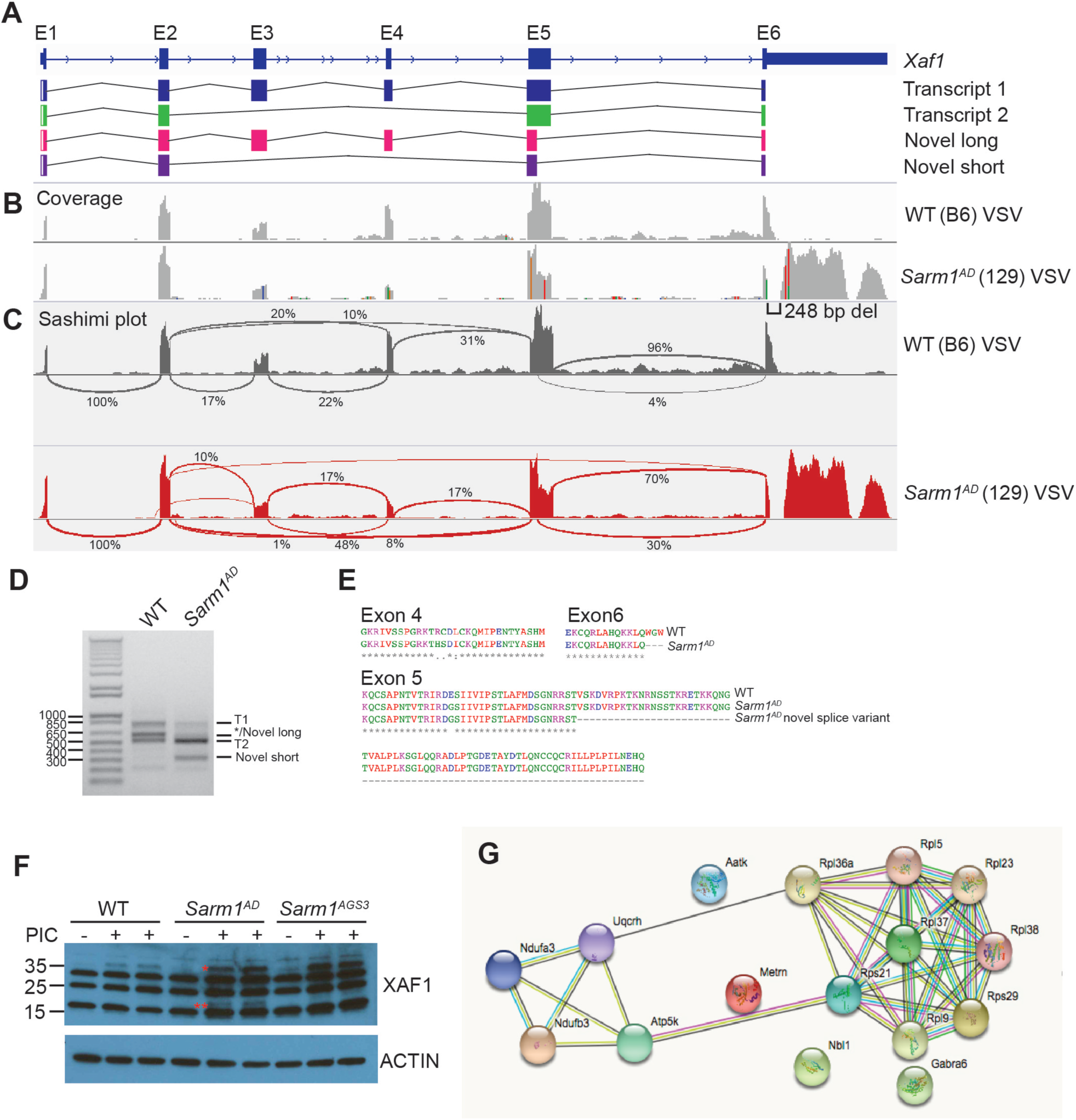
*Xaf1* sequence and isoform polymorphism. (**A**) *Xaf1* gene and transcripts. (**B**) WT and *Sarm1*^*AD*^ mice were infected with 10^7^ pfu of VSV and brain samples were collected for RNAseq at day 5 post-infection. Plots show coverage alignment of WT and *Sarm1*^*AD*^ sample reads to the B6 reference genome (mm10) at the *Xaf1* locus – colors indicate nucleotide changes from the reference sequence. (**C**) Sashimi plots (IGV) of the samples in B showing exon-exon splice junctions. (**D**) RT-PCR of *Xaf1* transcripts from samples in B. *Indicates that the ∼600 bp band corresponds to different transcripts in WT and *Sarm1*^*AD*^ samples. (**E**) Protein coding differences between WT (B6) and *Sarm1*^*AD*^ (129) *Xaf1* transcripts. (**F**) WT, *Sarm1*^*AD*^, and *Sarm1*^*AGS3*^ mice were injected i.v. with 100 ug of PIC, and splenocytes were isolated at 24 hrs for XAF1 western blot. *XAF1 isoform 1 * *Possible XAF1 novel isoform. (**G**) STRING analysis of significantly differentially expressed genes from mock-infected brainstem of WT and *Sarm1*^*AGS3*^ mice. Lines indicate known and predicted interactions.

Sashimi plots visualizing splice junctions showed an increase in junctions between exon 2 and 5 (10% to 48%) indicating less full length transcript in the *Sarm1*^*AD*^ strain, as well as a large increase in a novel splice variant between exon 5 and 6 (4% to 30%) in the *Sarm1*^*AD*^ strain (Fig 7C). Using RT-PCR primers directed against exon 1 and either the B6 or 129 exon 6, we detected the reported sequences for transcripts 1 and 2 in B6 (Fig 7D and see table III for sizes and accession numbers). In 129 we detected the reported sequence for transcript 1. The 3’ end of the 129 transcript 2 was incomplete in databases, and ended in the same sequence as transcript 1, resulting in the same C-terminal truncation. In the 129 samples we also detected two novel isoforms corresponding to the novel splice site between exon 5 and 6, leading to a novel long isoform (600 bp) similar to transcript 1 but lacking part of exon 5, and a novel short isoform (315 bp) similar to transcript 2 but also lacking part of exon 5. We detected a band of similar size to the novel long isoform in B6 (Fig 7E – indicated by *), however sequence analysis indicated this was a 626 bp transcript lacking exon 3 and leading to early truncation of the protein. The alternative splice site in exon 5 results in a large deletion of exon 5 (Fig 7E), but in-frame translation of exon 6. Importantly, the C-terminal domain is thought to be essential for binding to XIAP (31), and short isoforms are thought to function as dominant negative (22, 23), suggesting that these strain differences may lead to functional changes in XAF1.

**Table III.** *Xaf1* Transcripts

| Transcript | Size (bp) | Size (aa) | Size (kDa) | Accension |
| --- | --- | --- | --- | --- |
| B6 transcript 1 | 822 | 273 | 31 | ENSMUST00000146233.7 |
| 129 transcript 1 | 813 | 270 | 30 | MGP_129S1SvImJ_T0030476.1 |
| B6 transcript 2 | 537 | 178 | 20 | ENSMUST00000140842.8 |
| 129 transcript 2 | 528 | 175 | 20 | Submitted |
| 129 novel long | 600 | 199 | 23 | Submitted |
| 129 novel short | 315 | 104 | 12 | Submitted |

In order to test XAF1 antibodies, we generated XAF1-deficient 3T3 cell lines using CRISPR. Despite the presence of non-specific bands, using one of these antibodies we could detect XAF1 expression specifically in WT but not *Xaf1*^*-/-*^ cells (Fig S3). This band was only present following IFN treatment, in agreement with *Xaf1* being an interferon-stimulated gene. Importantly, the antibody epitope is present in all isoforms. Following treatment of mice with i.v. PIC to induce IFN, we were unable to detect XAF1 expression in the brain, but did observe expression in response to PIC treatment in the spleen. We observed a band corresponding to the size of the full-length protein in WT, *Sarm1*^*AD*^, and *Sarm1*^*AGS3*^ mice. However, we also observed a unique band in the *Sarm1*^*AD*^ strain following PIC treatment, which may represent either increased expression of isoform 2 or one of the novel isoforms (Fig 7F). No differences in Xaf1 expression levels were observed between WT and *Sarm1*^*AGS3*^ by RNAseq (Table IV), suggesting that SARM1 likely does not control XAF1 expression. Given the differential expression of XAF1 in the *Sarm1*^*AD*^ strain, and its known role in cell death, we speculate that XAF1 may account for some of the phenotypes described in this strain.

**Table IV.**
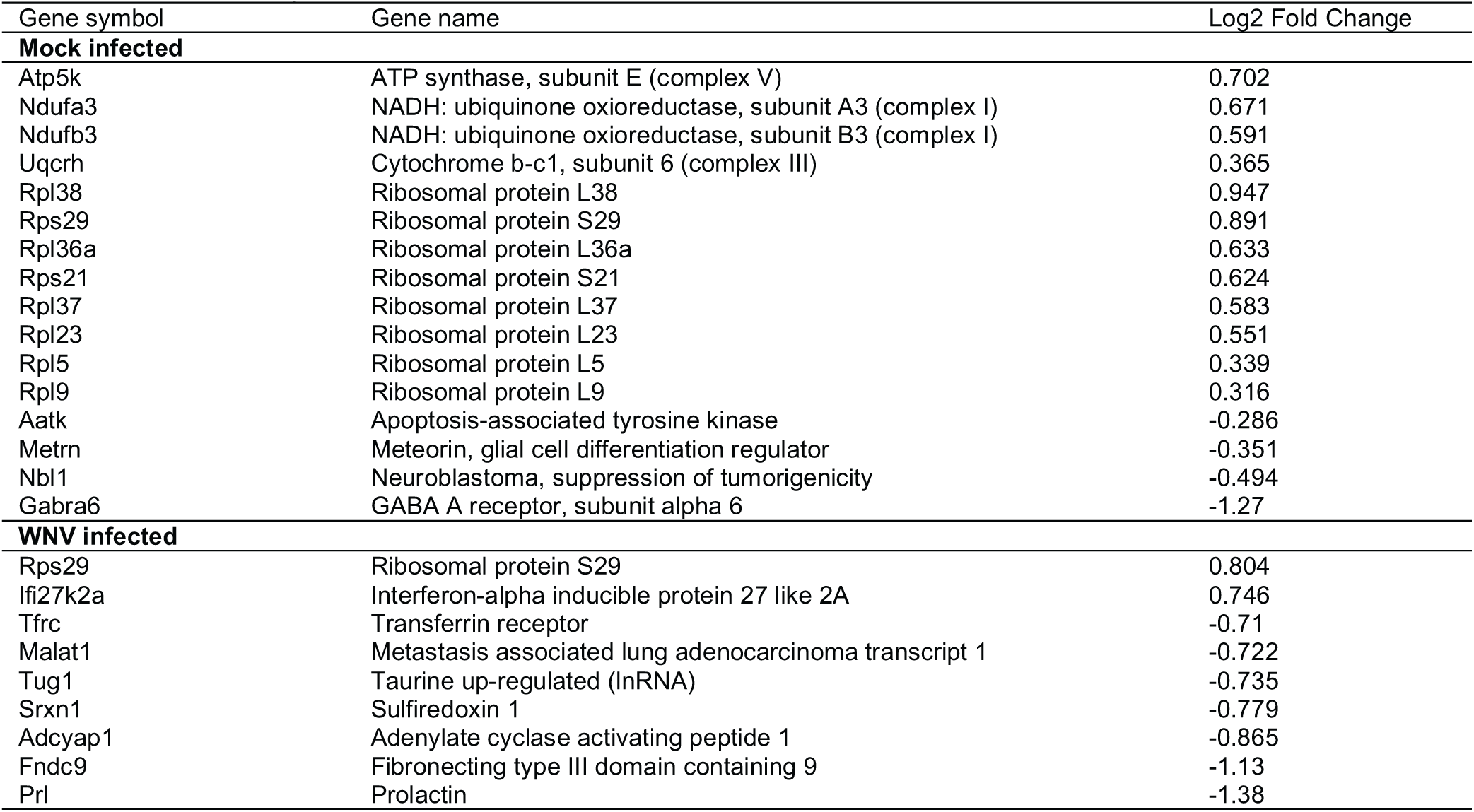
Differentially expressed transcripts between WT and *Sarm1*^*AGS3*^

### RNAseq on Sarm1 CRISPR mice

In order to understand possible functions for SARM1 we performed RNAseq on brainstem isolated from WT and *Sarm1*^*AGS3*^ mice infected with WNV or mock infected. In infected animals 9 transcripts were differentially regulated (Table IV). In mock infected animals 16 transcripts were differentially regulated, 4 of which are involved in the mitochondrial electron transport chain – Ndufa3 and Ndufb3 (complex I), Uqcrh (complex III), and Atp5k (complex V), as well as a number of small and large ribosomal proteins, and an apoptosis-associated tyrosine kinase (Table V and Fig 7D). In agreement with this data, a recent report suggests a role for SARM1 in mitochondrial respiration (8).

**Table V.**
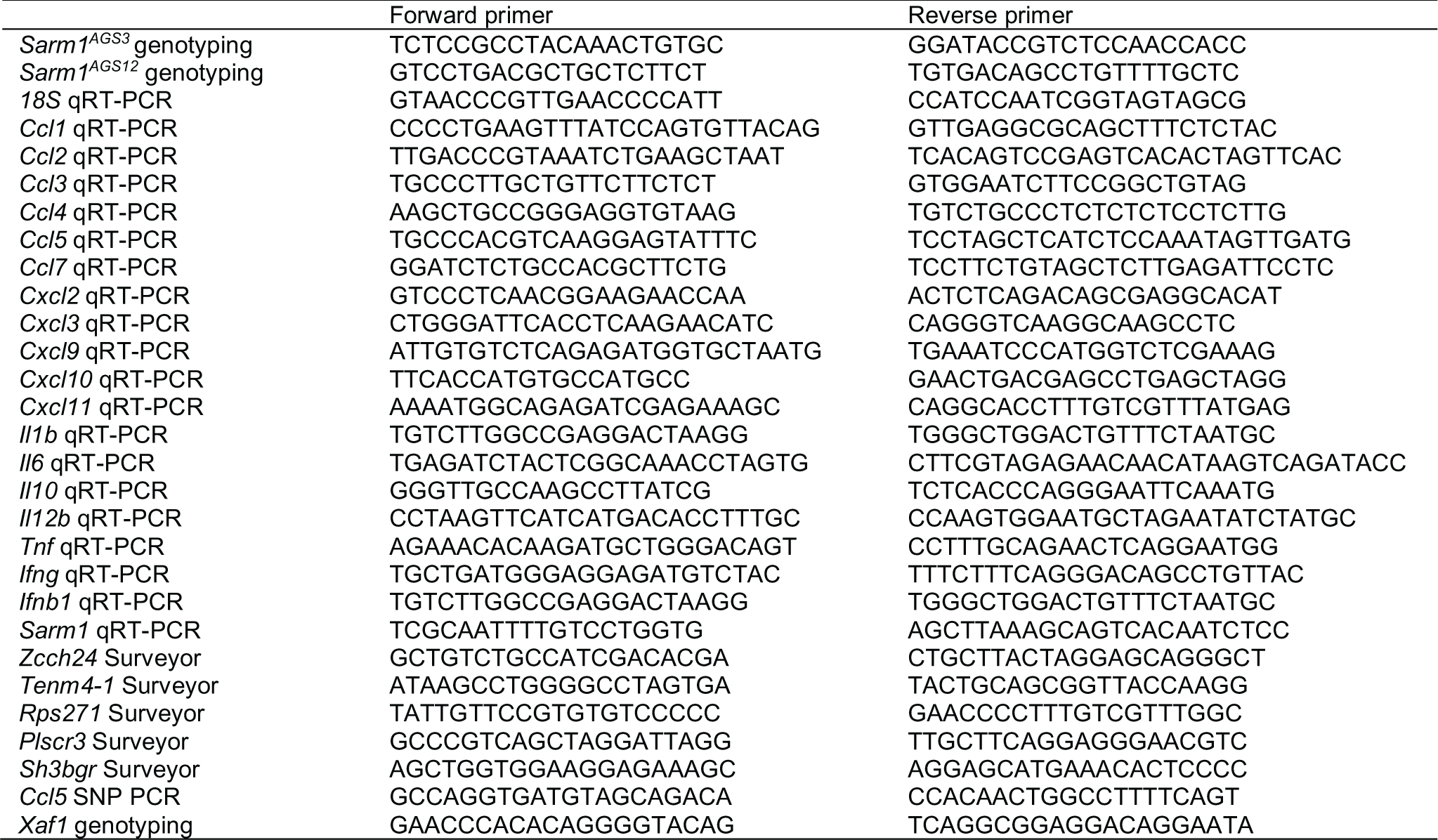
Primer sequences

## Discussion

Current evidence supports a role for SARM1 in axonal degeneration (2, 3). Roles for SARM1 in immunity have also been reported for CNS viral infections (15-17), but not for pathogens that replicate outside of the CNS including *M.tuberculosis, L. monocytogenes*, or influenza virus infection (17). Whether SARM1 plays a role outside of neural cells has proved difficult to answer. Studies on the expression and function of SARM1 have been hampered by the lack of reliable antibodies, making it difficult to gauge whether cells of the immune system express detectable protein levels. At the RNA level, evidence suggests predominant expression of SARM1 in the CNS. However, it remains possible that cells in the periphery express SARM1. We and others (15) did not detect the expression of SARM1 at the RNA level in macrophages, using primers that span exons 7 and 8, and detect high expression in WT but not *Sarm1*^*AD*^ brain. However, others report expression of a shorter 724 a.a. isoform in T cells and macrophages using primers spanning exons 5-7 (18, 32). Our primers should detect both isoforms, so the reason for the discrepancy is unclear.

In this study, we sought to address whether SARM1 plays a role in macrophages using cells from *Sarm1*^*AD*^ mice. Similar to published reports (18) we found differences in the production of *Ccl5*, as well as *Ccl3* and *Ccl4* in *Sarm1*^*AD*^ macrophages. However, a number of lines of evidence support that this is not due to SARM1 protein expression, but rather is due to background effects of the knockout strain. First, the defect in *Sarm1*^*AD*^ macrophages is limited to 3 particular chemokine genes that are located in close physical proximity to each other and the modified locus. Second, the defect is evident in response to a wide array of stimuli that induce different signaling pathways. Third, we could find no defects in the signaling components that are shared between the induction pathways for these stimuli. Fourth, siRNA knockdown failed to reproduce the *Sarm1*^*AD*^ chemokine phenotype suggesting a lack of dependence on SARM1 protein expression. Overexpression of SARM1 has been reported to modestly induce *Ccl5* expression (18), however we were unable to reproduce these findings. Additionally, we found differences in baseline expression of *Ccl3, Ccl4*, and *Ccl5* in unstimulated macrophages from *Sarm1*^*AD*^ mice, supporting an intrinsic difference. Finally, generation of new knockout strains on a pure genetic background also failed to support a role for SARM1 in macrophage chemokine production. These data in combination with the lack of expression/low expression of SARM1 in macrophages fail to support a role for SARM1 as a TLR adaptor protein in myeloid cells.

A variety of both protective and detrimental effects have been reported in different infection models in SARM1-deficient strains. These results are difficult to reconcile given the different construction of the knockout strains, and the significant variation in genetic background. Additionally, studies have not reported SNP analysis and whether or not additional backcrossing was done. SARM1 was reported to have a negative effect on susceptibility to both VSV and LACV infection, while it was reported to have a positive effect on susceptibility to WNV infection. We reported that *Sarm1*^*AD*^ mice were less susceptible to VSV, and showed lower cytokine responses and infiltration in the brain, while Mukherjee et al reported that *Sarm1*^*MSD*^ mice were protected from LACV infection, in a mechanism dependent on SARM1 interaction with MAVS (16). Our CRISPR knockout strains did not support a role for SARM1 in mediating this effect in either infection model. Surprisingly, none of our knockout lines - including *Sarm1*^*AD*^, *Sarm1*^*AGS3*^, *and Sarm1*^*AGS12*^ showed a protective effect during LACV infection, suggesting that the phenotype is specific to either the *Sarm1*^*MSD*^ strain or the viral strain. We found the *Sarm1*^*MSD*^ strain to differ from B6 at large portions of chromosome 10 and 11 in our analysis, which could account for the discrepant results. Additionally, the LACV original strain was used in our study, while Mukherjee et al used the LACV 1978 strain. These strains share 99% amino acid identity and are both highly virulent in young mice (33, 34), however differences in pathogenesis are observed in some strains (35). Our CRISPR knockout strains did, however support a role for SARM1 in mediating the positive effect during WNV infection. Surprisingly, the *Sarm1*^*AD*^ line showed similar susceptibility to WT mice during WNV infection. Both the *Sarm1*^*AD*^ and *Sarm1*^*MSD*^ lines were made on the 129 background, however the *Sarm1*^*AD*^ line retains neomycin. Similar phenotypes in *Sarm1*^*AGS3*^, *Sarm1*^*AGS12*^, and *Sarm1*^*MSD*^ mice suggest that either neomycin effects on neighboring genes, or other 129 background effects account for the different phenotype of the *Sarm1*^*AD*^ *s*train to WNV.

Here we show background strain-dependent differences in the expression of the proapoptotic protein XAF1, which may represent a good candidate gene for the protective effect described in the knockout strains, however a number of other possibilities are consistent with the data. The protective phenotype could be due to: 1) differences in chemokine levels due to the 129 congenic locus, which can also influence immune cell infiltration 2) transcriptional interference from neomycin effecting chemokines or other neighboring genes within the congenic interval 3) other mutations within the congenic interval or 4) other background effects. We had originally reported that *Sarm1*^*AD*^ mice had lower levels of monocyte and macrophage infiltration into the brain, in agreement with their lower cytokine/chemokine levels, and postulated that this may lead to protection from immune-mediated tissue damage (17). Neomycin has been documented to abrogate downstream gene expression and interfere with locus control regions at both short and at megabase distances (36-38), which would also be consistent with lower recruitment of Pol II to the *Ccl5* promoter in *Sarm1*^*AD*^ mice (18). In addition, the importance of genetic background on the phenotype of knockout mice is well known – and examples of interfering passenger mutations abound in the literature (39).

This example and others highlight the advantages of generating new knockout strains using CRISPR technology. Given the difficulty in accessing the expression of SARM1 using available antibodies, we have used this approach along with a homology-directed repair template to generate mice with epitope-tagged SARM1. This line was unfortunately lost, however similar approaches will be important for assessing SARM1-interacting proteins and signaling pathways *in vivo*. RNAseq in our CRISPR strains suggests loss of SARM1 expression leads to changes in expression of ribosomal, and mitochondrial electron transport chain genes. This is in agreement with a recent study showing that SARM1 phosphorylation regulates NAD+ cleavage leading to inhibition of mitochondrial respiration (8). Overall the data suggest that reevaluation of phenotypes described in SARM1-deficient strains will be important for understanding the function of SARM1 in different contexts.

## Acknowledgements

This work was partially supported by R01AI108715 to JKL. The mice used for this study were produced by the Mouse Genetics and Gene Targeting Center of Research Excellence (CoRE), which is supported by the Icahn School of Medicine at Mount Sinai, and a Cancer Center Support Grant (1P30CA196521-01) from the National Cancer Institute/National Institutes of Health. We thank William Janssen at the Microscopy CoRE and Advanced Bioimaging Center for assistance with nerve imaging. We thank Michael Diamond and Andrew Pekosz for reagents, and Zuleyma Peralta and Maryline Panis for technical assistance.

## Materials and Methods

### Mice

*Sarm1*^*AD*^ mice on the C57BL/6J background were generated previously from 129 ES cells (1) and backcrossed to C57BL/6J 10 generations. Mice were compared to WT C57BL/6J mice purchased from Jackson. Animal studies were approved by the Institutional Animal Care and Use Committee of Icahn School of Medicine at Mount Sinai. CRISPR knockout mice were generated using the CRISPR design tool to select the guide sequence TCGCGAAGTGTCGCCCGGAGTGG in exon 1 of the *Sarm1* gene. This was cloned into pSpCas9(BB)-2A-GFP (Addgene) as described (29). The resulting plasmid was injected at 1 ng/ul into the male pronuclei of one-cell stage C57BL/6J mouse embryos. After injection, the embryos were returned to the oviducts of pseudopregnant Swiss-Webster (SW) females that had been mated the day before with vasectomized SW males. Resulting pups were characterized using a combination of PCR, sequencing, and surveyor analysis. *Sarm1*^*AGS3*^ were genotyped by PCR using the primers listed in table I and the PCR conditions 95° 30 sec, 53° 30 sec, 72° 1 min. *Sarm1*^*AGS12*^ were genotyped by PCR using the primers listed in table I and cycling conditions 95° 30 sec, 63.5° 30 sec, 72° 1 min, and sequencing using the forward primer. Surveyor assay was performed using *Sarm1*^*AG3S*^ PCR conditions and Surveyor Mutation Detection Kit (IDT) followed by electrophoresis on Novex 20% TBE gels (Invitrogen). Off-target CRISPR cleavage was accessed by PCR amplification using the primers listed in table I and cycling conditions 95° 30 sec, 60° 30 sec, 72° 1 min and the Surveyor Mutation Detection Kit (IDT) on a pup from a cross of the *Sarm1*^*AGS3*^ founder mouse to WT.

### SNP analysis

To determine the precise genetic background of *Sarm1*^*AD*^ and *Sarm1*^*MSD*^ mice, 384 SNP panel analysis was performed by Charles River Genetic Testing Services. Testing was performed on tail DNA from *Sarm1*^*AD*^ mice maintained in our colony and MEF DNA derived from the *Sarm1*^*MSD*^ line (provided by Michael Diamond) because the Diamond lab no longer maintains the animal colony. *Ccl5* SNPs were genotyped by PCR of genomic DNA from C57BL/6J or *Sarm1*^*AD*^ mice using primers listed in table I and cycling conditions 95° 30 sec, 60° 30 sec, 72° 1 min, followed by cloning into pGEM-T (Promega) and sequencing.

### Macrophages and 3T3 cell lines

Bone marrow was obtained from femurs and tibias of mice, RBCs were lysed and cells were cultured for 7 days in RPMI 1640 (Gibco) containing 10% FBS (Hyclone), Penicillin, Streptomycin, L-glutamine, Hepes (Cellgro), β-ME, and 10 ng/ml rmM-CSF (R&D Systems). Macrophages were removed from the plate following incubation with cold PBS and plated in 24-well plates at 0.25×10^6^/well. Cells were stimulated the following day with Poly(I:C) HMW (Invivogen), *E.coli* 0111:B4 LPS purified by gel filtration (Sigma), R848 (Invivogen), CL075 (Invivogen), NDV-GFP (40), or VSV Indiana strain at concentrations listed in figure legends. *3T3 Xaf1*^*-/-*^ cells were generated by cloning the guide sequences AGCTTCCTGCAGTGCTTCTGTGG and AGGCTGACTTCCAAGTGTGCAGG located in exon 1 of Xaf1 into pSpCas9n(BB)-2A-GFP, and transfecting into 3T3 cells using LTX (Invitrogen). Single cell clones were obtained by limiting dilution and screened by PCR using the primers listed in table I and the PCR conditions 95° 30 sec, 60.2° 30 sec, 72° 30 sec, Surveyor assay (as above), and western blot.

### qRT-PCR

Total RNA was extracted from macrophage cultures using EZNA total RNA kit and RNase-free DNase (Omega). RNA was reverse-transcribed using Maxima Reverse Transcriptase and oligo-dT (Thermo). Quantitative RT-PCR was performed on cDNA using LightCycler 480 SYBR Green I Master Mix (Roche) and the primers listed in table I on a LightCycler 480 II. Data is shown as relative expression (2^-ΔΔCt^ relative to 18S).

### ELISAs

CCL3 ELISA was performed using the mouse CCL3/MIP-1α DuoSet (R&D Systems), TNF-α ELISA was performed using the mouse TNF ELISA kit (BD OptEIA), and IFN-α ELISA was performed using the Verikine Mouse IFN Alpha ELISA Kit (PBL Assay Science) according to the manufacturer’s instructions.

### Western blots

For macrophage blots, 0.5×10^6^ macrophages were plated in 12-well plates. The following day cells were serum starved for 3 hrs, stimulated with 10 ng/ml LPS or TNF-α for the indicated amount of time, lysed in RIPA buffer containing Halt Protease and Phosphatase Inhibitor Cocktail (Thermo), denatured in Laemmli buffer, run on 4-12% Bis-Tris gels (Invitrogen), and transferred to PVDF membranes. For Xaf1 blots 8×10^3^ 3T3 cells were treated for 24 hrs with 2000 U universal type I IFN (PBL) and lysed in Laemmli buffer. Mice were injected with 100 ug of HMW Poly(I:C) in 200 uL of PBS, spleens were harvested at 24 hrs. and homogenized in RIPA containing cOmplete Protease Inhibitor Cocktail (Roche), denatured in Laemmli buffer, run on 4-12% Mini-Protean gels (BioRad), and rapid transferred to PVDF membranes. Membranes were blocked with 0.2% I-BLOCK (Applied Biosystems) 0.1% Tween-20 in TBS and probed with rabbit IκBα (Cell Signaling 9242), rabbit phosphor-SAPK/JNK (Thr183/Tyr185) (Cell Signaling 9251), rabbit phosphor-p44/42 MAPK (Erk1/2) (Thr202/Tyr204) (Cell Signaling 4370), rabbit phosphor-p38 MAPK (Thr180/Tyr182) (Cell Signaling 9211), and mouse phosphor-Akt (Ser 473) (587F11) (Cell Signaling 4051), and rabbit Xaf1 (aa166-194, LS Bio LS-C158287), followed by detection with ECL donkey anti-rabbit IgG HRP or ECL sheep anti-mouse IgG HRP (GE Healthcare), or directly detected with rabbit β-Actin HRP (Cell Signaling 5125), or mouse V5-HRP (Serotec).

### Ca^2+^ signaling

Macrophages were plated at 0.75×10^5^/well in 96-well black clear-bottom plates overnight. Cells were loaded with 10 μM Fura-2-AM in 0.1% BSA in Hanks buffer for 30 min, washed, and fluorescence was measured (330 nm->513 nm – 380 nm->513) on a plate reader after addition of 1 mM ATP or 0.1 ug/ml LPS.

### RAW-SARM1-V5 cells and siRNA

Full length *Sarm1* with a C-terminal V5 tag or the V5 tag alone was cloned into the pLVX-IRES-Puro lentiviral vector (Clontech) and transfected into 293T cells along with gag/pol and VSV-G expression plasmids to generate lentiviral particles. These were used to infect RAW 264.7 cells, followed by puromycin selection. Expression was checked by western blot and immunofluorescence. SARM1 was knocked down using Dharmacon Accell siRNA targeting *Sarm1* (target sequences: UGCUGUUGCUCGAUUCGUC and CCAAGGUGUUCAGCGACAU). 0.3×10^5^ RAW-V5 or RAW-SARM1-V5 cells were plated in 96-well plates, the following day siRNA was added at 1 μM in Accell delivery media for 72 hrs, Accell delivery media was removed and DMEM containing 10% FBS was added for 3 hrs. Cells were stimulated with 10 ng/ml LPS for 3 hrs and qPCR was performed as above. Knockdown in primary macrophages was performed similarly on 0.5×10^5^ cells.

### VSV, LACV, and WNV infection

6-8-week old female mice were anesthetized with ketamine/xylazine and infected intranasally with 10^7^ pfu of VSV Indiana strain in 20 μl PBS. Mice were monitored daily for weight and sacrificed when exhibiting severe paralysis or more than 25% weight loss. For brain cytokines, mice were perfused with PBS and brains were removed and stored in RNAlater, followed by homogenization and RNA isolation with EZNA HP Total RNA kit (Omega), and qPCR as above. In BSL3 containment, 8-week old female mice were anesthetized with isoflurane and injected subcutaneously in the neck with 10^2^ FFU of West Nile virus-NY99 in 50 ul of PBS and monitored as for VSV. 3-week old male and female mice were infected intraperitoneally with 10^3^ pfu of LACV original (parent) strain (kindly provided by Andrew Pekosz) and sacrificed when exhibiting severe paralysis.

### RNAseq

WT and *Sarm1*^*AD*^ mice were infected intranasally with 10^7^ pfu of VSV and brain RNA was prepared as above at day 5 post-infection. RNA quality and quantity was assessed using the Agilent Bioanalyzer and Qubit RNA Broad Range Assay kit (Thermo Fisher), respectively. Barcoded directional RNA-Sequencing libraries were prepared using the TruSeq Stranded Total RNA Sample Preparation kit (Illumina). Libraries were pooled and sequenced on the Illumina HiSeq platform in a 100 bp single-end read run format. After adapter removal with cutadapt (https://doi.org/10.14806/ej.17.1.200) and base quality trimming to remove 3′ read sequences if more than 20 bases with Q≥20 were present, paired-end reads were mapped to the murine mm10 reference genome using STAR (v2.5.3a) (41) and reference gene annotations from ENSEMBL (v75). WT and *Sarm1*^*AGS3*^ mice were infected intracranially with 100 FFU of WNV-Kunjin strain in 30 ul PBS or mock infected with PBS. Animals were perfused with PBS at day 5 post-infection. RNA preparation and sequencing was performed as above except that sequencing was non-directional and used a NextSeq machine with 150 bp reads. Protein-protein association networks were determined using STRING database (42). RNAseq datasets have been deposited in GEO under the record numbers GSE136221 and GSE136284, and *Xaf1* transcripts have been deposited in Genbank under the accession number (submitted).

### Sciatic nerve transections

WT and *Sarm1*^*AGS3*^ were anesthetized with ketamine/xylazine, fur was shaved, and skin was cleaned. An incision was made in the skin and the muscle was separated to expose the sciatic nerve. A 1 mm portion of the nerve was excised, and the skin was closed with staples. Antibiotic ointment was applied to the incision and 0.05 mg/kg buprenorphine was administered immediately and at 6 hrs for pain. Mice were housed for 14 days, and euthanized with 15% aqueous choral hydrate, followed by perfusion with 1% Paraformaldehyde/PBS, pH 7.2 at a flow rate of 7.5 mls/min, and immediately with 2% paraformaldehyde and 2% glutaraldehyde/PBS, pH 7.2 at the same flow rate for an additional 10 minutes. Skin was removed, and the carcass placed in immersion fixation (same as above) to be post-fixed for a minimum of one week at 4 degrees C. The transected and non-transected nerves were removed and flat mold embedded to ensure cross-sectional orientation in EPON resin. Polymerized blocks were sectioned on a Leica UC7 Ultramicrotome using a histoknife at 0.5 um, counterstained with 1% Toluidine Blue and coverslipped. Brightfield images were acquired with an Axioimager Z2M microscope (Zeiss) with an EC Plan-Neofluar 40x/1.3 oil objective, and processed with Fiji software (NIH).

**Figure S2.**
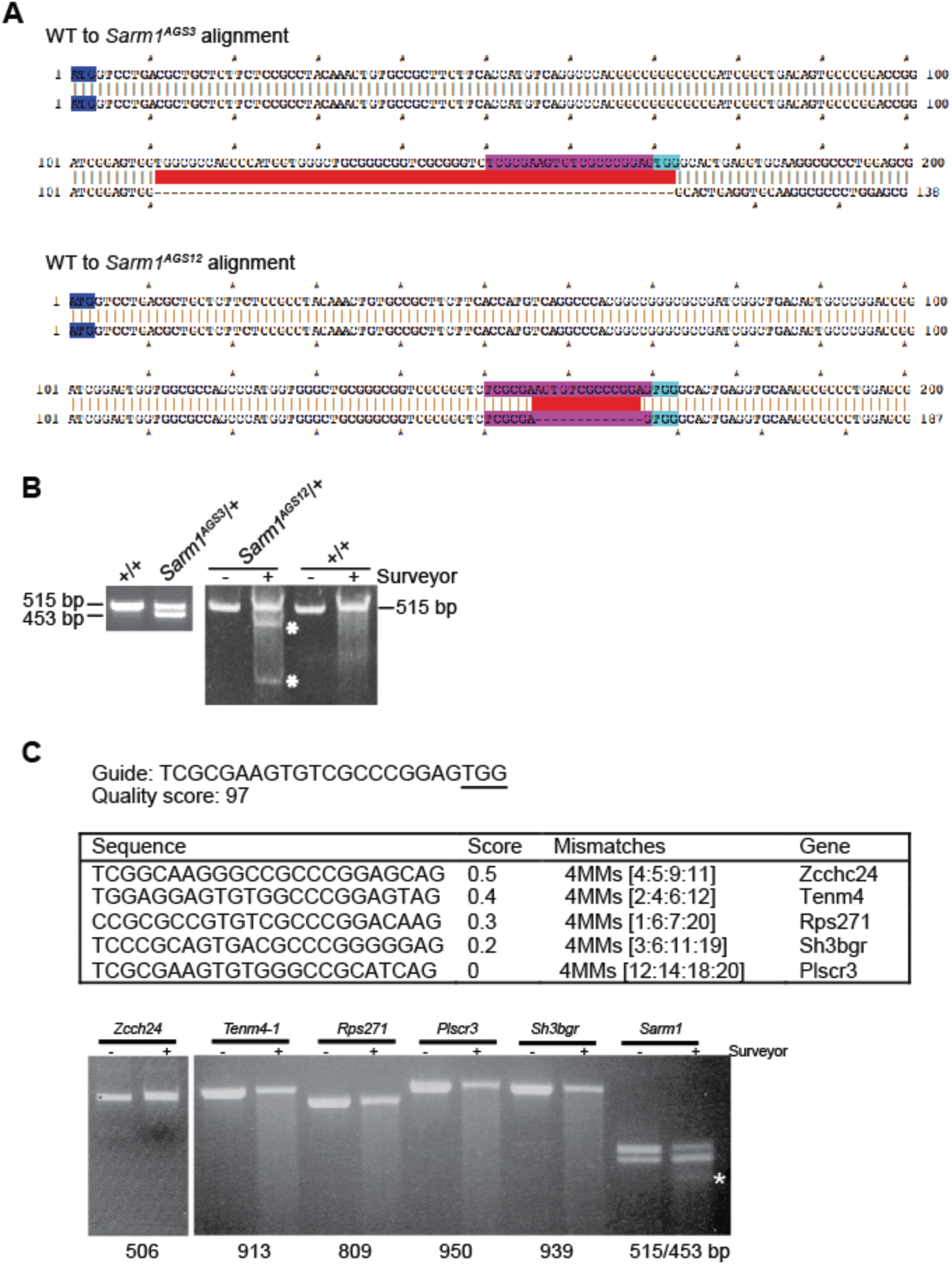
SARM1 CRISPR alleles and the absence of off-target CRISPR-mediated cleavage in *Sarm1*^*AGS3*^ mice. (A) Sequence alignment of WT and *Sarm1*^*AGS3*^, and WT and *Sarm1*^*AGS12*^ alleles. The start codon is indicated in blue, the guide sequence in pink, and the PAM sequence in cyan. (B) WT or *Sarm1*^*AGS3*^/+ genomic DNA was amplified by PCR and run on an agarose gel (left). WT or *Sarm1*^*AGS12*^/+ genomic DNA was amplified by PCR, digested with Surveyor Nuclease, and run on a TBE gel (right). (C) Genes with off-target cleavage sites in exonic regions were amplified by PCR of genomic DNA from a cross of the *Sarm1*^*AGS3*^ founder mouse to WT, digested with Surveyor Nucelase, and run on agarose gels. Sizes of PCR products are listed below the gel. The Sarm1 locus served as a positive control for Surveyor cleavage. Note both the *Sarm1*^*AGS3*^ and *Sarm1*^*AGS12*^ alleles resulted from the same mosaic founder mouse.

**FIGURE S3.**
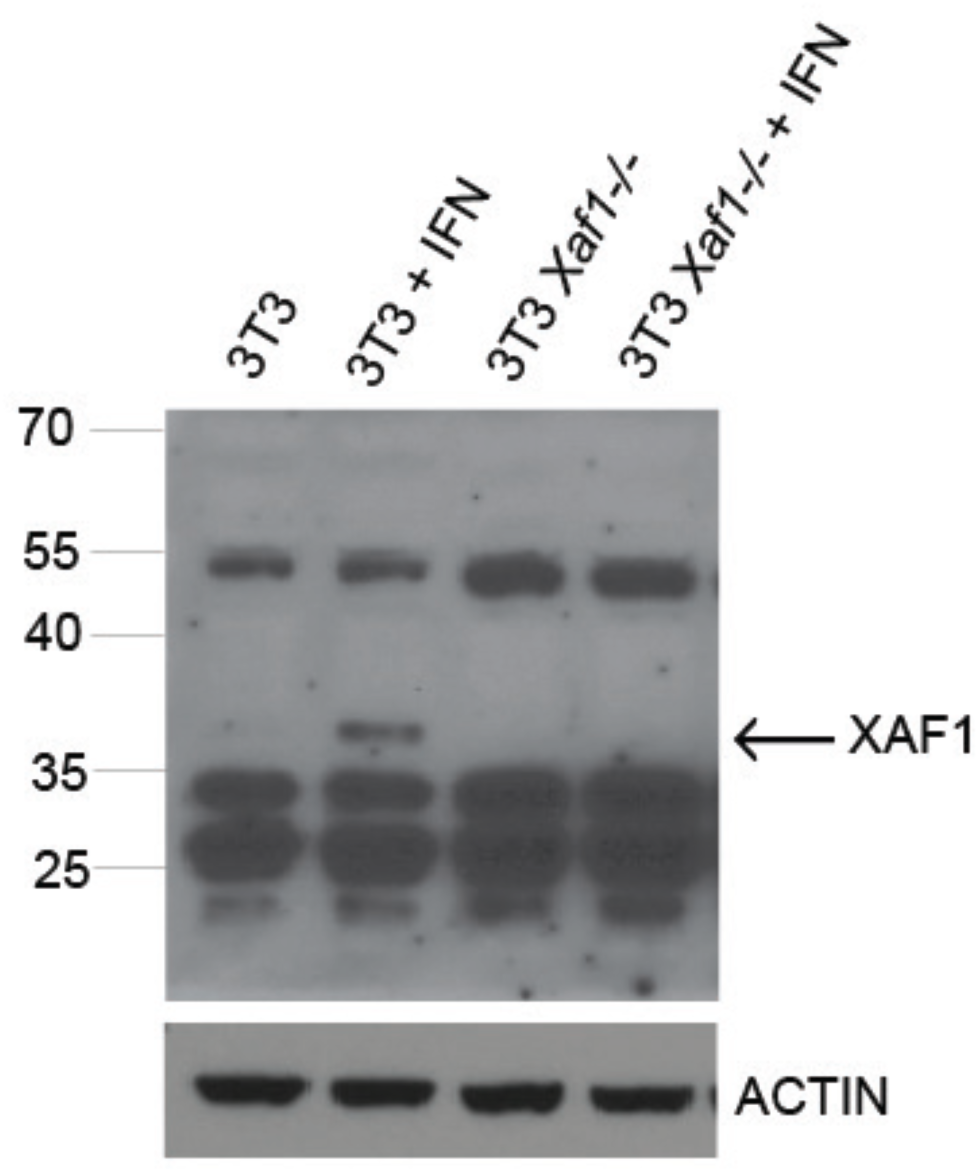
Specificity of XAF1 antibody. 3T3 and 3T3 *Xaf1*^*-/-*^ cells were mock treated, or treated with 1000 U of universal type I IFN for 24 hrs before western blotting for XAF1.

